# Metassembler: Merging and optimizing de novo genome assemblies

**DOI:** 10.1101/016352

**Authors:** Alejandro Hernandez Wences, Michael C. Schatz

## Abstract

Genome assembly projects typically run multiple algorithms in an attempt to find the single best assembly, although those assemblies often have complementary, if untapped, strengths and weaknesses. We present our metassembler algorithm that merges multiple assemblies of a genome into a single superior sequence. We apply it to the four genomes from the Assemblathon competitions and show it consistently and substantially improves the contiguity and quality of each assembly. We also develop guidelines for metassembly by systematically evaluating 120 permutations of merging the top 5 assemblies of the first Assemblathon competition. The software is open-source at http://metassembler.sourceforge.net.

## Rationale

Next generation high-throughput DNA sequencing technologies are being used to tackle an increasing list of biological questions [1]. One of the most fundamental uses is for *de novo* genome assembly, where the goal is to reconstruct the genome sequence of an organism from high throughput sequencing data, while dealing with their characteristic short reads and error rates [2]. Genome assembly is fundamental to computational biology, as a successful assembly is needed to study the gene content, regulatory regions, or evolutionary relationships in a genome, along with several other questions. As such, it is critical that researchers can create the best possible assembly from the available data.

While de novo genome assembly has been studied for more than twenty years, the problem is far from being solved. The available assembly algorithms vary most significantly in the techniques and heuristics applied to assemble repetitive sequences and resolve errors present, especially in response to the ever-changing landscape of available biotechnologies [2-4]. The central challenge in genome assembly is repetitive sequences can give rise to false or ambiguous overlaps, leading to the termination of contigs and/or the introduction of errors [5]. Indeed, all assemblers can assemble non-repetitive error-free data with ease.

As a result, the performance of different *de novo* genome assembly algorithms can vary greatly on the same dataset, although it has been repeatedly demonstrated that no single assembler is optimal in every possible quality metric [6-8]. The most widely used metrics for evaluating an assembly include 1) contiguity statistics such as scaffold and contig N50 size, 2) accuracy statistics such as the number of structural errors found when compared to an available reference genome (GAGE evaluation tool [8]), 3) presence of core eukaryotic genes (CEGMA [9]) or, if available, transcript mapping rates, and 4) the concordance of the sequence with remapped pair-end and mate-pair reads (REAPR [10], assembly validation [11], or assembly likelihood [12]).

The performance of different assemblers, as measured by these metrics, has recently been systematically compared in the two international Assemblathon competitions [6, 7], as well as other evaluations, where different researchers generated the best possible assemblies of the same sample using different algorithms and parameters. The first Assemblathon competition used a simulated genome derived from a mutated version of human chromosome 13, thus enabling evaluation directly with the truth. The second Assemblathon competition used real data from the three species: a Fish (*Maylandia zebra*), a Bird (*Melopsittacus undulatus*), and a Snake (*Boa constrictor constrictor*), whose complete reference genomes are unavailable and therefore relied on reference-free assembly evaluation metrics. Dozens of assemblies were generated for each species, and the results of both competitions show that the different algorithms have varying strengths and weaknesses; that is, a single assembly may maximize a subset of evaluation metrics but no single assembly or assembler maximized all the metrics at once in every dataset. These projects even demonstrated that different parameter settings of a single algorithm could significantly vary the results.

To overcome this challenge and make best use of the available algorithms and data, we present our Metassembly algorithm for merging and optimizing multiple assemblies together into a single superior assembly. The metassembly combines the locally best sequence from all input assemblies at each region of the genome, and merges them into a final sequence as good as or superior to the constituent assemblies. The merging is performed with an iterative, progressive approach where the current metassembled sequence is aligned and revised pairwise with each available assembly. After aligning the current metassembly sequence with the next assembly, it evaluates any conflicts and selects the locally best sequence as assessed by the compression-expansion (CE) statistic proposed by Zimin et al [13]. Unlike previous works including the assembly reconciliation algorithm [13] and GAM-NGS [14], our approach works with current high throughput sequence data and is designed to merge multiple assemblies all together. Our algorithm also has the capability to close more types of scaffolding gaps, and a scaffolding function whenever the alignment information and the local mate-pair reads support such modifications.

We tested our algorithm in the four genomes of the Assemblathon competitions, and demonstrate marked improvement in the contiguity and accuracy of each. A critical aspect to our merging algorithms is determining the order in which the input assemblies should be evaluated. To address this question, we systematically computed the metassembly of all possible 120 permutations of the top five assemblies of the Assemblathon 1 competition. Our algorithm achieved an average improvement of 4.6Mb for scaffold N50 size while improving or maintaining quality statistics such as the number of duplicated sequences and genomic rearrangements. For the Assemblathon 2 competition, we metassembled the top six assemblies for each of the three species available. Similar improvements were obtained by significantly increasing contiguity statistics (contig and scaffold N50 size) while maintaining overall quality. These results show the compelling nature of the metassembly algorithm for all future genome assembly projects, and we have released the software and documentation to the community open-source at http://metassembler.sourceforge.net.

## Metassembler Algorithm

The metassembly algorithm performs pairwise, progressive alignments to merge multiple assemblies in the order specified by the user or by some metric. The algorithm makes no assumptions on the way the input assemblies are generated, and can be applied to a set of assemblies generated with different software packages, parameters and/or different types of data. The only data requirement is at least one jumping library is available to evaluate the presence of compression/expansion mis-assemblies, although that data type need not been used in any or all of the assemblies.

### Pairwise merging process

The pairwise merging process follows the basic logic of the assembly reconciliation algorithm, and uses one of the input assemblies as the “primary” assembly and the other as the “secondary” assembly that is used to add information to the primary; in particular, it is used to correct insertion/deletion errors, close gaps, and to scaffold sequences in the primary assembly.

**Figure 1:**
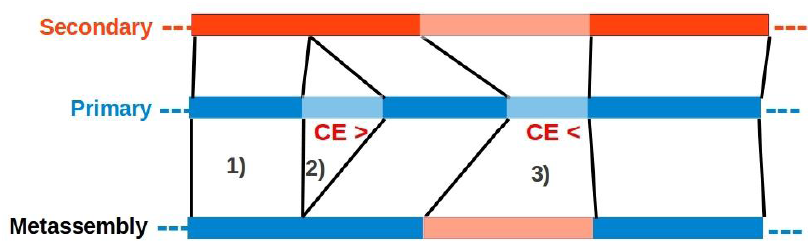
Schematic representation of the pairwise merging process. Dark color represents alignment blocks between the primary and secondary assemblies. Light color represents unaligned sequences. 1) For blocks of aligned sequence, the algorithm inserts the primary sequence to the new Metassembly. 2) Insertion in the primary with respect to the secondary assembly: because the CE statistic is a large positive value (>3) for the primary sequence, the algorithm skips the primary insertion and chooses the secondary sequence instead. 3) Both assemblies have a unaligned insertion: because the primary insertion is shorter than the secondary insertion, and because the primary has a large negative CE statistic (<-3), the algorithm will choose the secondary insertion over the primary, thus correcting the CE statistic.

The pairwise merging process consist of 4 steps:

1. **Whole Genome Alignment:** The metassembler algorithm uses the *nucmer* program from the *MUMmer* package to align the two input assemblies [15]. The resulting alignments are then refined using *delta-filter (also part of MUMmer)*, to compute the 1-to-1 mapping between the primary and secondary assemblies. As a result, each position in the primary sequence will be uniquely mapped to the best corresponding position in the secondary as assessed by a LIS (Longest Increasing Subsequence) maximization function weighted by the product of the length and identity of the alignment. This optimization step identifies the most significant correspondence between the two assemblies, discarding any repeat induced spurious alignment. By default our algorithm only takes into account those scaffolds present in the filtered set of alignments, i.e. it discards any sequences (scaffolds and/or contigs) with zero significant alignments from both assemblies. This generally eliminates spurious sequences that are most commonly short error-prone sequences. However, the user can optionally retain all sequences with a minimum length regardless of whether they have any significant alignments if so desired.
2. **Assembly evaluation:** The algorithm maps the mate pair libraries to both the primary and secondary assemblies using the short read mapping algorithm Bowtie2 [16]. The algorithm uses the resulting insert lengths to compute the CE statistic [13] at every base pair using a plane-sweep approach. The CE-statistic quantifies how compressed or expanded the set of mate pairs spanning a given position are in comparison to the expected insert size. Formally, it computes a z-test to detect statistically significant differences between the local mean insert size and the global (expected) mean insert size. Values substantially less than zero (typically < −3) indicate high probability of compression, and values substantially greater than zero (typically > +3) indicate high probability of expansion. These values are used by the merging step to assess which assembly is more likely to be correct in the event of a conflicting alignment. It is worth noting that even though there may be outlying correctly assembled regions of the genome with high absolute value of the CE statistic (erroneously signaling for a misassembly), the metric is not used as an error detection statistic in isolation, but rather as a method to locally compare candidate sequences.
3. **Assembly comparison and merging:** The metassembly algorithm scans each primary sequence to identify segments of aligned and unaligned sequences indicating gaps or discrepancies. Every aligned segment of the primary sequence is automatically added to the metassembly; in contrast, when a difference is found, the algorithm compares the CE statistic and coverage at the corresponding breakpoint positions to determine which of the two sequences will be added to the metassembly sequence (**Figure 1**).

For insertion/deletion events, the algorithm replaces the primary assembly sequence with the corresponding sequence of the secondary if all the following conditions are met:

1. ***Poor primary assembly:** abs(CE primary) > z*, where abs(CE primary) is the absolute value of the CE statistic in the primary assembly at the breakpoint position, and ‘*z*’ is a user specified threshold (3 by default).
2. ***Improved secondary assembly:** abs(CE primary) – abs(CE secondary) > d*, where ‘*d’* is a user specified threshold (2 by default).
3. ***Improved metassembly:*** If the insert coverage in the primary sequence is greater than zero then we can infer the CE statistic value resulting from choosing the secondary sequence instead of the primary. To do this we first compute an estimate of the local mean insert length after the modification: *y*_*i*_^*^ = *Y*_*i*_ - *primary insertion/deletion length + secondary insertion/deletion length*, where Yi is the primary assembly mean insert length. We then use *Y*_*i*_^*^ to compute the inferred CE statistic after the modification (*CE*_*i*_^*^). The algorithm only makes changes to the primary if *abs*(*CE*_*i*_^*^) < *z*.
4. ***Improved metassembly:*** The algorithm only makes changes to the primary assembly if *abs(CE secondary) < z*

A “scaffold gap” in the primary assembly is defined as a segment with at least *t* contiguous gap bases (N’s) such that *p* percent of the entire segment is gap sequence. Both, *t* and *p* are parameters (*t* = 50, *p* = 0.65 by default), allowing isolated gap nucleotides to be skipped and neighboring gaps to be joined into a single unit. In contrast, gaps in the secondary assembly are defined as having at least 10 contiguous gap bases and at least 10% of gap sequence to maximize sensitivity. If a gaps in the primary assembly is spanned by a non-gap sequence in the secondary assembly, the primary sequence is replaced by the secondary if the improved metassembly conditions 3) and 4) above are met.

It is often the case that secondary sequences do not entirely align to a particular primary sequence, having “overhangs” at the very ends of the sequence caused by errors or lower coverage at the very end of the contig. These cases are also handled as gap closure or insertion/deletion events.

**4) Scaffolding:** Our algorithm also finds the set of primary sequences that can be linked into scaffolds. If two primary sequences align contiguously to the same secondary scaffold after filtering for repetitive alignments, and the secondary sequence has coverage above a user specified threshold (*default:20*), then the two primary sequences are linked together into a single scaffold or a single contig, depending on if the secondary sequence has a scaffold gap at that position.

### Progressive analysis

After the pairwise merging has been completed with the top two assemblies, the algorithm iterates the procedure using that newly formed metassembly and the next best assembly as inputs (**Figure 2**). Assemblies are processed according to the user specified ordering or ranking scheme, such as ordering by assembly contiguity (N50 size, etc) or completeness metrics (CEGMA, etc). For example, in our analysis below we have found that ranking assemblies from largest to smallest by their contig N50 size is generally effective heuristic.

**Figure 2:**
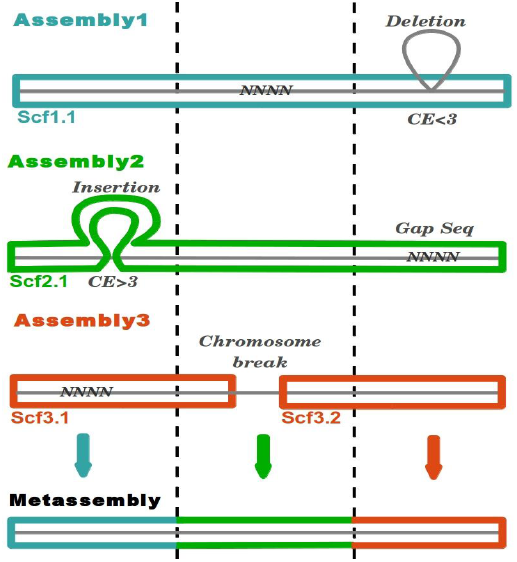
Schematic diagram of the progressive metassembly of 3 assemblies. All three input assemblies have gap sequences and a variety of errors such such that no pair of assemblies will create a perfect assembly. However, the final metassembly of all 3 assemblies together will reconstruct the entire correct genome.

The progressive pairwise merging avoids the computational load of performing whole genome alignments for all pairs of assemblies. Moreover, because the overall contiguity of the intermediate metassembly increases with each merging step, the subsequent whole genome alignment and mate-pair read alignments become more sensitive; thus improving error detection, CE statistic accuracy, and the overall performance of the following pairwise merge.

## Evaluation

We applied our metassembly algorithm to four genomes with multiple publicly available assemblies: the Asssemblathon1 competition genome, and the three genomes (Snake, Bird, and Fish) of the Assemblathon2 competition. For each dataset, we compared overall contiguity statistics such as the N50 size and total span at each merging step, the change in CE-statistic at positions were merges were made, and various accuracy metrics depending or not if a reference genome was available.

### 1. Metassembly of the Assemblathon 1 Genome

The Assemblathon 1 genome consists of a pair of simulated haploid genomes generated with the Evolver evolution tool (http://www.drive5.com/evolver/) using the human chromosome 13 sequence and annotation as input. Paired-end and mate-pair reads were then simulated from the resulting 112Mbp genome, which were then presented as an international competition to create the best possible assembly of the data. More than 40 entries were submitted and evaluated by an ensemble of quality metrics. For our analysis, we focused on the top five assemblies reported by the Assemblathon1 overall rank shown in Supplemental Table 1.

We systematically metassembled all 120 possible permutations of the five input assemblies, using the 2.5kbp mate-library to evaluate the CE status of each assembly (**Figure 3,** Supplementary Table 1, and Supplementary Note 2). Because the genome has an exact reference available, we were able to compute 10 different quality metrics including the number of major structural errors and the corrected scaffold and contig N50 sizes using the GAGE assembly evaluation tool. The corrected scaffold N50 size and corrected contig N50 size are computed by splitting the input sequences at places where significant errors are found relatively to the reference assembly, and then computing the N50 sizes of the remaining sequences.

**Figure 3:**
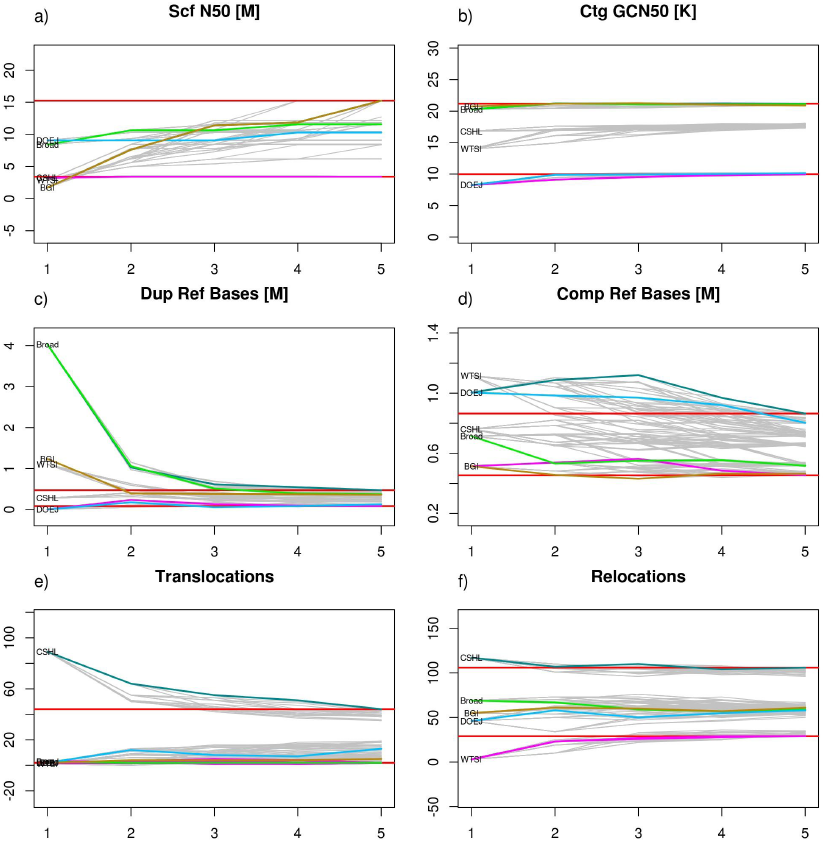
Assemblathon 1 Metassembly Accuracy. Assembly contiguity and accuracy metrics are plotted at each merging step for all possible permutations of the five input assemblies: A) Scaffold N50, B) Corrected contig N50, C) Duplicated reference bases, D) Deleted reference bases, E) Translocations, and D) Relocations. For all plots, the x-axis represents the number of input assemblies being metassembled, with 1 being the starting assembly. The two horizontal red lines mark the final maximum and minimum value of the metric across all permutations. Most of the permutations are plotted in gray, while permutations of particular note are plotted with different colors: The pink line represents the permutation that has the maximum value in the final metassembly while the dark blue line represents the permutation with the minimum value. Also, the green line represents the permutation resulting from ordering the input assemblies by the overall rank reported in the Assemblathon 1 paper (Broad-BGI-WTSI-DOEJGI-CSHL), the light blue line represents the permutation obtained by ordering the input assemblies by scaffold N50 size (DOEJGI-Broad-WTSI-CSHL-BGI) while the brown line represents the order by contig N50 size (BGI-Broad-CSHL-WTSI-DOEJGI).

On average 138.9 scaffold merges were made across all permutations, as well as 2288 gaps and 541 indels were processed (maximum of 5899 and 1178 respectively). As shown in **Figure 3a**(and Sup Fig 1a), scaffold NG50 sizes (N50 sizes relative to the reference genome size) consistently increase using any permutation of the input assemblies: the mean difference in scaffold NG50 size between the final metassembly and the starting assembly is 4.6 Mb, with a maximum improvement of 13.5 Mb for the BGI-Broad-CSHL-DOEJGI-WTSI permutation (starting with the BGI assembly as the primary assembly and adding the remaining assemblies in that order). Contig NG50 sizes also improve substantially with a mean increment of 17.3Kb and a maximum of 70Kb (Sup Fig 1c). Furthermore, GAGE corrected scaffold NG50 size (Scf GC-NG50, Sup Fig 1b) and corrected contig NG50 size (Ctg GC-NG50, **Figure 3b**) are also substantially increased with a mean difference between the final metassembly and the initial assembly of 701Kb and 1.5Kb respectively. We also evaluated the change in CE-statistic at positions where modifications to the primary assembly were made and find the vast majority of events have a positive difference. This indicates that the CE-statistic is reduced closer to zero, further supporting that our algorithm is capable of correcting such events without introducing errors. (Sup. Figure 3)

These results imply that during the metassembly process, the scaffolds are becoming much larger and more contiguous without sacrificing contig or scaffold quality. The other accuracy metrics further support this conclusion: the number of duplicated reference bases and the number of deleted reference bases (**Figures 3c and 3d**) significantly decrease with a mean difference of −1Mb and −460Kb respectively, while the number of translocations (mean difference: −5.4) and relocations (mean difference: 4) do not show any significant change. The complete table of metrics for all metassemblies and their corresponding boxplots are included in the Supplemental Material.

### 2. Metassembly ordering

We studied the dependency between the order in which the assemblies are metassembled and the quality of the final metassembly by evaluating the final metassembly of all input permutations. To do so, we computed the Z-score of assembly quality, proposed in the Assemblathon 2 paper, which aggregates and summarizes all of the different metrics into a single value based on the mean and standard deviation of the individual metrics. The boxplots shown in **Figure 4** summarize the distribution of overall Z-scores for all the metassemblies starting with each of the input assemblies. This shows that our algorithm is capable of significantly improving Overall Z-scores with a mean increment of 14.5 standard deviations, but also strongly suggests that quality and contiguity of the final assembly is dependent on the order of merging and which assembly is used first.

**Figure 4.**
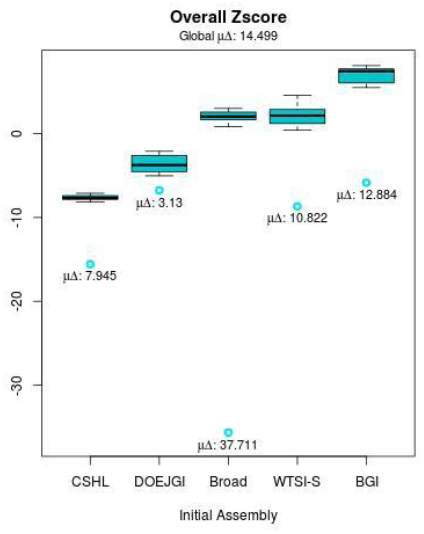
Boxplots of overall Z-scores for metassemblies grouped by initial assembly. Blue circles indicate the Z-score of the corresponding initial assembly. Below each circle, the corresponding mean difference in Z-scores between the final metassembly and the initial assembly (µ) is shown. The global mean difference is also shown at the top.

This dependency could be inflated if the quality metrics were redundant or highly correlated, so we also evaluated the distribution of overall Z-scores using just the subset of statistically independent metrics obtained by an Independent Component Analysis (ICA) to select the most statistically independent metrics (Supplemental Note 3 and 4). The ICA-selected subset of statistically most informative metrics were: 1) Inversions, 2) Compressed Reference Bases, 3) Missing Reference Bases, and 4) Relocations using the top 50% components in the kurtosis distribution, plus 5) Duplicated Reference Bases for the top 80%. Using just these subsets of quality metrics, the same dependency between initial assembly and final Overall Z-score was observed (Sup Figure 2).

Given that the final metassembly quality is dependent on the order in which the input assemblies are processed, we considered whether there was a simple ordering rule that would lead to the best (or nearly best) metassembly as measured by the overall Z-score. We therefore correlated reference-independent metrics of the initial input assemblies, such as scaffold N50 size and contig N50 size, with the median value of their corresponding overall Z-score distribution. We found that contig N50 size correlates positively with median overall Z-score (r=0.72 and permutation test p-value=0.08), while scaffold N50 does not correlate (r= −0.22). These results were reproduced when evaluating just the subset of metrics selected by the ICA, contig N50 size had a correlation of r=0.65 with pval=0.12, while scaffold N50 size had a correlation of r= −0.24. This shows that ordering the initial assemblies by contig N50 size should give a high quality metassembly. Indeed, the BGI-Broad-CSHL-WTSI-DOEJGI permutation (ordered by contig N50 size) falls in position 13 of the total 120 ranked metassemblies, while the permutation DOEJGI-Broad-WTSI-CSHL-BGI (ordering by scaffold N50 size) lies in position 85. The permutation ordered by Assemblathon 1 rank (Broad-BGI-WTSI-DOEJGI-CSHL) lies at position 39.

### 3. Metassembly of the three Assemblathon 2 genomes

For each of three species in the Assemblathon 2 project we applied our algorithm to the top 6 assemblies as ranked by the cumulative Z-score reported in the paper (Supp. Note 2). For the Fish genome, we excluded the top ranking CSHL entry and picked the lesser ranking CSHL/ALLPATHS-LG based assembly instead, since the top ranking CSHL entry used a prototype of our metassembly algorithm. Since these genomes are much larger than the Assemblathon1 genome and because we had already established strong rules for ordering the assemblies, we generated three metassemblies for each species: ordering the input assemblies by contig N50 size, scaffold N50 size, or Assemblathon 2 cumulative Z-score (A2Z) (Sup Table 3).

We evaluated the correctness and contiguity of the metassembly at each merging step using the metrics used by the Assemblathon 2 evaluation. Namely, we evaluated the presence of core eukaryotic genes using the CEGMA algorithm [9], as well as the concordance of the metassembly sequence with remapped pair-end and mate-pair reads using REAPR [10]. The former looks for the presence of 248 highly conserved core eukaryotic genes in the assembly sequence as a proxy for the completeness and accuracy of the assembly, especially of genes. The latter evaluates errors by aligning paired-end and mate-pair libraries and looking for regions where coverage drops or the distribution of observed fragment lengths differs from the expected distribution. It then splits the assembled sequence at places where errors are found to compute corrected scaffold N50 size (Scf RC-N50) and corrected contig N50 sizes (Ctg RC-N50). In our analysis we recomputed these relative to the estimated genome size (Scf RC-NG50 and Ctg RC-NG50 respectively). We also evaluated the CE statistic at positions where modifications to the starting assembly were made (Supplementary Figure 4).

In all three species, the contiguity statistics are significantly improved by our metassembly algorithm (**Figure 5**). Contig NG50 sizes increased by at least 3.9Kb and 4.4Kb for the Fish and Snake metassemblies, with a maximum increment of 3.98Kb and 13.8Kb respectively. The largest increment in Contig NG50 size was observed in the Bird species, with an improvement ranging between 43.9Kb and 69.1Kb. Moreover, scaffold NG50 sizes improved by between 0.96Mb and 1.4Mb for the Snake genome, and between 105Kb and 122Kb for the Fish genome. For the Bird species a decrement of −3.9Kb is observed for the A2Z permutation, while a maximum increment of 1.9Mb is observed for the Scf N50 order permutation. Furthermore, the assembly quality metrics either remain unchanged or show a tendency to improve. The REAPR corrected NG50 sizes increase throughout the metassembly process, as well as the percentage of error free bases. The number of CEGMA genes found either increases (in the Bird and Snake assemblies) or decreases slightly because of poor secondary assemblies (in the Fish genome). The complete results are available as supplementary files.

**Figure 5.**
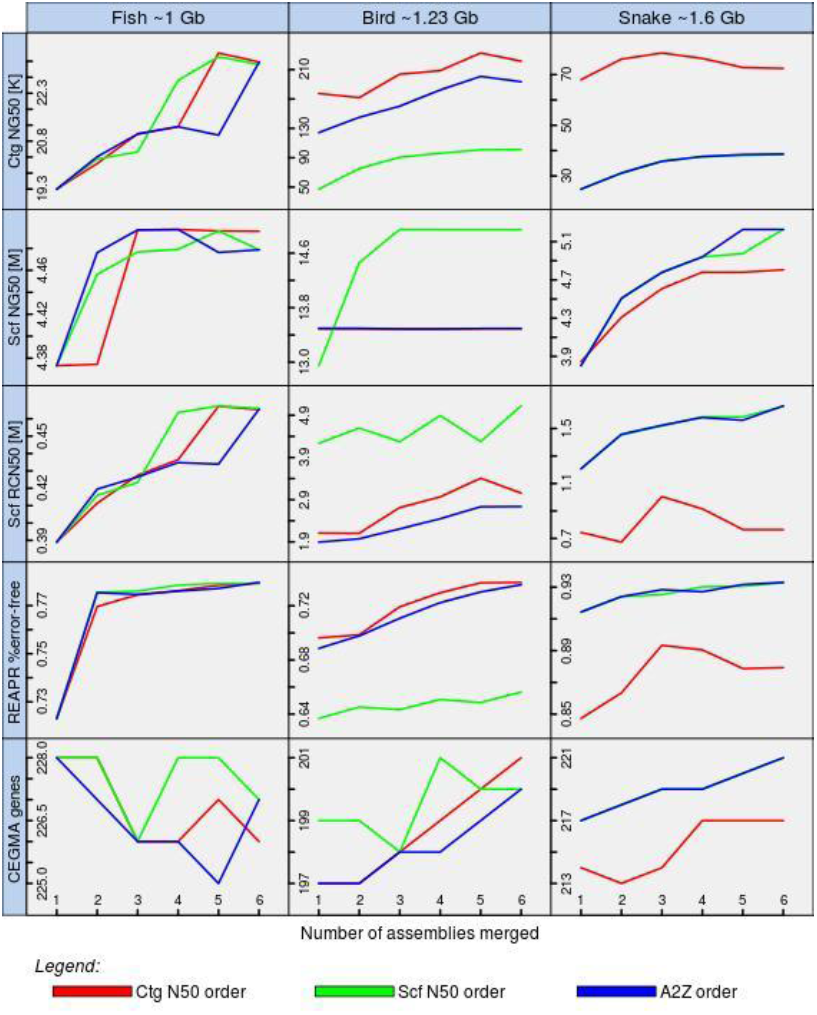
Assemblathon 2 Metassembly contiguity and accuracy metrics. Assembly contiguity and accuracy metrics are shown at each merging step of all metassemblies for the three species. The x axis represents the number of assemblies merged with one being the initial input assembly.

These results show that our algorithm is capable of improving assembly contiguity and quality metrics with real sequence data. The Fish genome shows the least improvement in scaffold N50 size probably because the secondary assemblies provide little new scaffolding information: The BCM Fish genome assembly has the largest scaffold N50 size, and was the starting assembly for all three permutations attempted. Furthermore, the number of CEGMA genes found for the Fish genome slightly decreases (from 228 to 225), due to the sequence filtering done at the alignment step. The effects of this filtering could have been exacerbated by the inclusion the SGA assembly that has a scaffold N50 size of only 0.1Mb compared to 1.24Mb for the second lowest scaffold N50 size. For the other genomes, CEGMA results improve through the metassembly process, especially when the metassemblies are computed in order of the contig N50 sizes.

We also ranked the input assemblies and metassemblies by overall Z-score using the following metrics: Scaffold NG50 size, Contig NG50szie, Scaffold REAPR-Corrected NG50 size, Contig REAPR-Corrected NG50 size, Percentage of error free bases, and number of CEGMA genes found. In the Fish and Snake species the three metassemblies occupy the top three positions, while in the Bird species the three metassemblies lie within the top four positions.

## Discussion

We have developed an algorithm capable of merging multiple assemblies generated with different algorithms, parameters, and possibly different sequence data into a single metassembly. We demonstrated the power of our algorithm in both simulated and real data over 4 species, 23 assemblies, and more than one hundred metassemblies. Given that the final metassembly depends critically on the order in which the input assemblies are incorporated we computed the metassembly for all 120 possible permutations to systematically explore this relationship. We found that scaffold NG50 size is improved for all permutations with a mean increase of 4.6 Mb while corrected scaffold and corrected contig NG50 sizes improve by 701Kb and 1.5Kb on average. For the Assemblathon 1 and 2 datasets we show that scaffold NG50 and contig NG50 sizes are substantially improved in all species while quality statistics remain practically unchanged or improve. These results support that our algorithm is capable of improving contiguity statistics without any loss in sequence accuracy; in fact a prototype version of this algorithm was used for our Assemblathon 2 Fish submission, and was one of the highest rated algorithms in the entire competition. New ranking of the assemblathon genomes show our version wins or is extremely competitive in all 4 species.

These observations together indicate that combining information from multiple assemblies into a single assembly is a powerful approach towards improving assembly quality and contiguity prior to its publication or subsequent analysis. Furthermore these results show our open-source algorithm is capable of performing such a task in a fast and accurate way. Indeed, consensus approaches, such as metassembly, are often a highly effective strategy for optimizing complex decisions, as long as the underlying algorithms can utilize independent characteristics [17]. In the case of genome assembly, the metassembly algorithm is synthesizing the various heuristics and techniques implemented for error correction and repeat resolution in each of the assemblers used. In this light, future work remains to develop additional error correction and repeat resolution modules that can systematically explore the range of possible algorithms for them, although care must be taken to keep the search space tractable.

## Abbreviations

CEGMA -: Core Eukaryotic Genes Mapping Approach
CE-statistic -: Compression/Expansion statistic
Comp Ref Bases -: Compressed Reference Bases
Ctg NG50 -: Contig N50 size relative to the estimated/reference genome size
Ctg GC-NG50 -: Contig GAGE Corrected N50, relative to the reference genome size Ctg RC-NG50 - Contig REAPR Corrected N50, relative to the estimated genome size Dup Ref Bases - Duplicated Reference Bases
GAGE -: Genome Assembly Gold Standard Evaluation
GAM-NGS -: Genomic Assemblies Merger for Next Generation Sequencing
ICA -: Independent Component Analysis
PCA -: Principal Components Analysis
REAPR -: Recognising Errors in Assemblies using Paired Reads
Scf NG50 -: Scaffold N50 size relative to the estimated/reference genome size
Scf GC-NG50 -: Scaffold GAGE Corrected N50, relative to the reference genome size
Scf RC-NG50 -: Scaffold REAPR Corrected N50, relative to the estimated genome size

## Competing Interests

The author(s) declare that they have no competing interests.

## Authors contributions

MCS designed the study. AHW implemented the software and performed the experiments. Both authors wrote and approved the manuscript.

## Acknowledgments

We would like to thank Paul Baranay and Scott Emrich for their helpful discussions and involvement during the development of the prototype of the software. The project was supported in part by National Institutes of Health award (R01-HG006677) and by National Science Foundation awards (DBI-1350041 and IOS- 1237880) to MCS.

